# De novo design of high-affinity protein binders to bioactive helical peptides

**DOI:** 10.1101/2022.12.10.519862

**Authors:** Susana Vázquez Torres, Philip J. Y. Leung, Isaac D. Lutz, Preetham Venkatesh, Joseph L. Watson, Fabian Hink, Huu-Hien Huynh, Andy Hsien-Wei Yeh, David Juergens, Nathaniel R. Bennett, Andrew N. Hoofnagle, Eric Huang, Michael J MacCoss, Marc Expòsit, Gyu Rie Lee, Paul M. Levine, Xinting Li, Mila Lamb, Elif Nihal Korkmaz, Jeff Nivala, Lance Stewart, Joseph M. Rogers, David Baker

## Abstract

Many peptide hormones form an alpha-helix upon binding their receptors^1–4^, and sensitive detection methods for them could contribute to better clinical management. *De novo* protein design can now generate binders with high affinity and specificity to structured proteins^5,6^. However, the design of interactions between proteins and short helical peptides is an unmet challenge. Here, we describe parametric generation and deep learning-based methods for designing proteins to address this challenge. We show that with the RF*diffusion* generative model, picomolar affinity binders can be generated to helical peptide targets either by noising and then denoising lower affinity designs generated with other methods, or completely *de novo* starting from random noise distributions; to our knowledge these are the highest affinity designed binding proteins against any protein or small molecule target generated directly by computation without any experimental optimization. The RF*diffusion* designs enable the enrichment of parathyroid hormone or other bioactive peptides in human plasma and subsequent detection by mass spectrometry, and bioluminescence-based protein biosensors. Capture reagents for bioactive helical peptides generated using the methods described here could aid in the improved diagnosis and therapeutic management of human diseases.^7,8^

## Main

Peptide hormones, such as parathyroid hormone (PTH), neuropeptide Y (NPY), glucagon (GCG), and secretin (SCT), which adopt alpha helical structures upon binding their receptors^1–4^, play key roles in human biology and are well established biomarkers in clinical care and biomedical research (Fig. 1a). There is considerable interest in their sensitive and specific quantification, which currently relies on antibodies that require substantial resources to generate, can be difficult to produce with high affinity, and often have less-than-desirable stability and reproducibility^5^. Furthermore, the loop-mediated interaction surfaces of antibodies are not particularly well suited to high specificity binding of extended helical peptides. Designed proteins can be readily produced with high yield and low cost in *E. coli* and have very high stability, but while there have been considerable advances in *de novo* protein design to generate binders for folded proteins^5,6^, the design of proteins that bind helical peptides with high affinity and specificity remains an outstanding challenge. Design of peptide-binding proteins is challenging for two reasons. First, proteins designed to bind folded proteins, such as picomolar affinity hyper-stable 50-65 residue minibinders^5^, have shapes suitable for binding rigid concave targets, but not for cradling extended peptides. Second, peptides have fewer residues to interact with, and are often partially or entirely unstructured in isolation^9^; as a result, there can be an entropic cost of structuring the peptide into a specific conformation^10^, which compromises the favorable free energy of association. Progress has been made in designing peptides that bind to extended beta strand structures^11^ and polyproline II conformations conformations^12^ using protein side chains to interact with the peptide backbone, but such interactions cannot be made with alpha helical peptides due to the extensive internal backbone - backbone hydrogen bonding.

**Figure 1.**
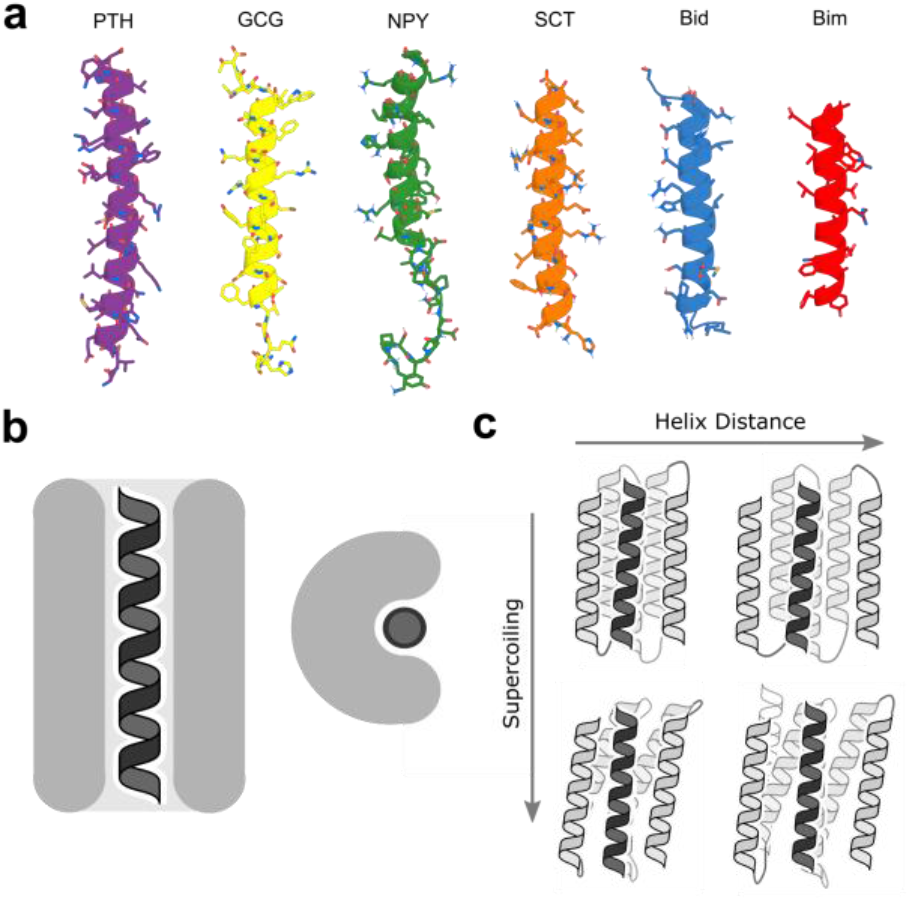
Binding helical peptides in groove scaffolds. **(a)** Helical peptide targets: parathyroid hormone (PTH), glucagon (GCG), neuropeptide Y (NPY), secretin (SCT), and the apoptosis-related BH3 domains of Bid and Bim. **(b)** “Open groove” structural solution to the helix binding problem. **(c)** Parametric approach to sampling of groove scaffolds varying supercoiling and helix distance to fit different targets.

### Design of helical peptide binding scaffolds

We set out to develop general methods for designing proteins that bind peptides in helical conformations. To fully leverage recent advances in protein design, we explored both parametric and deep learning-based approaches. For parametric generation, we reasoned that helical bundle scaffolds with an open groove for a helical peptide could provide a general solution to the helical peptide binding problem: the extended interaction surface between the full length of the helical peptide target and the contacting helices on the designed scaffold could enable the design of high affinity and specificity binding (Fig. 1b). In parallel, we reasoned that deep learning methods, which do not pre-specify scaffold geometries, could permit the exploration of different potential solutions to helical peptide binding.

### Parametric design of groove scaffolds

We began by exploring parametric methods for generating backbones with overall “groove” shapes. Using the Crick parameterization of alpha-helical coiled coils^13^, we devised a method to sample scaffolds consisting of a three-helix groove supported by two buttressing helices (Fig. 1c, see Supplementary Materials). We assembled a library of these scaffolds sampling a range of supercoiling and helix-helix spacings to accommodate a variety of helical peptide targets (Supplementary Fig. S1). We then used this library to design binders to PTH, GCG, and NPY, and screened 12 designs for each target using a nanoBiT split luciferase binding assay. Many of the designs bound their targets (3/12, 4/12, and 8/12 to PTH, GCG, and NPY) but with only micromolar affinities (see Supplementary Materials). These results suggest that groove-shaped scaffolds can be designed to bind helical peptides, but also that design method improvement was necessary to achieve high-affinity binding.

While powerful for generating and sampling a large number of potential scaffolds, the parametric generation approach has the limitation of building only from ideal building blocks, in this case parametric alpha helices. Deep learning methods do not have these limitations, and we explored whether RoseTTAFold inpainting (RF*joint*)^14^, a model that can jointly design protein sequences and structures, could be used to improve the modest affinities of our parametrically-designed PTH binders (Fig. 2a). We used RF inpainting to extend the binders (non-parametrically) to incorporate additional interactions with the target peptide to take advantage of the full potential binding interface of the peptide. Out of 192 designs tested, 44 showed binding against PTH in initial yeast display screening. Following SEC purification, the best binder was found to bind at 6.1 nM affinity to PTH. Binding was quite specific: very little binding was observed to PTH related peptide (PTHrp), a related peptide sequence with 34% sequence identity (Fig. 2A). Overall, the affinity of the starting PTH binders was improved by approximately three orders of magnitude, and the highest-affinity binder had 19% greater surface area contacting the target peptide. We used the same design strategy to generate higher affinity binders for NPY and GCG. Using weak parametric binders as a starting point, we extended their binding interfaces and generated a ∼231 nM affinity binder for GCG and a 3.5 μM binder for NPY after screening 96 designs (Supplementary Fig. S2).

**Figure 2.**
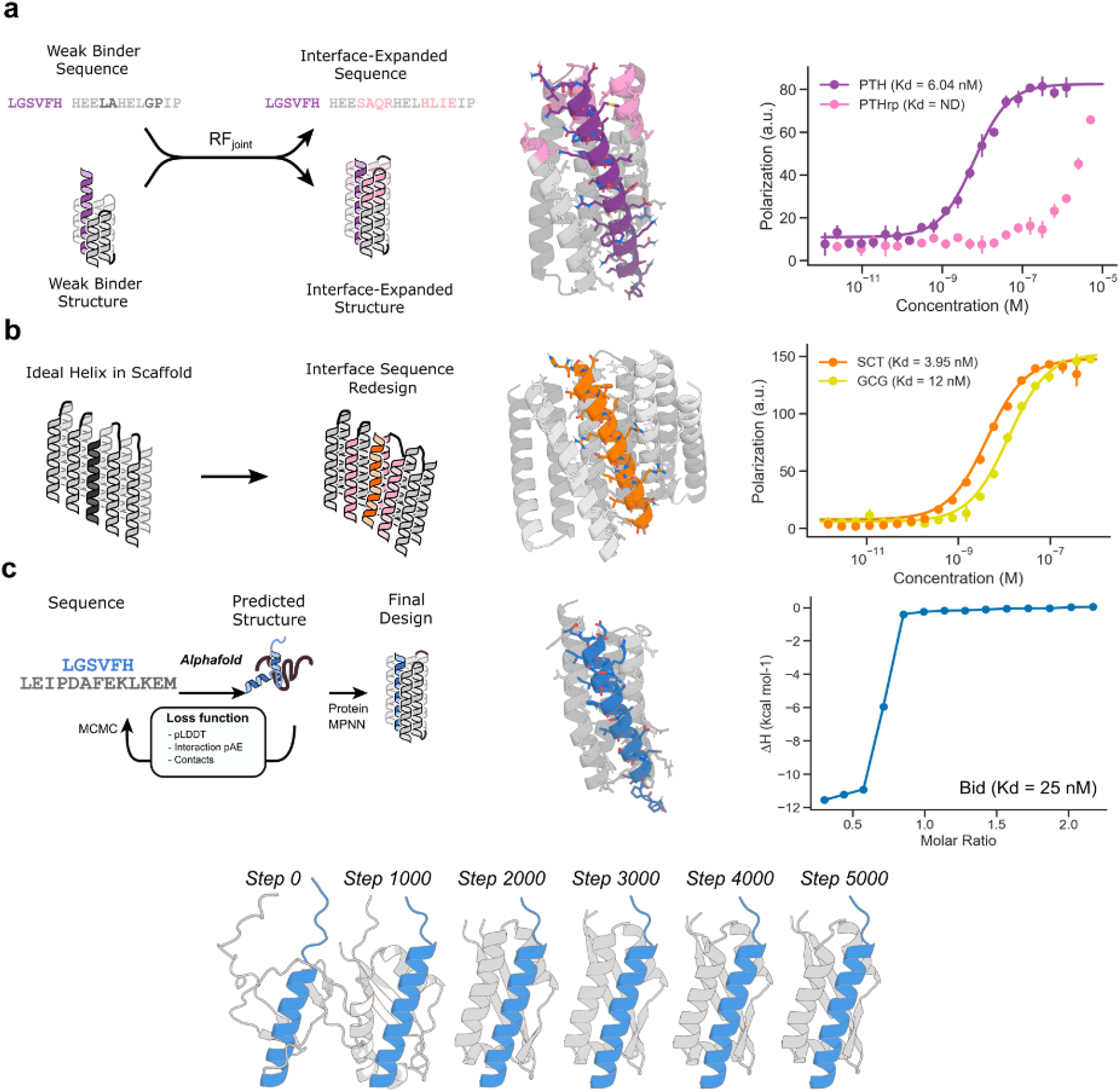
Design strategies for binding helical peptides. **(a)** Inpainting binder optimization: redesign of parametrically generated binder designs using RF*joint* inpainting to expand the binding interface. Left: schematic illustration of approach. Middle: original parametric scaffold (gray), inpainted design with extended interface (pink), and PTH target (purple). Right: Fluorescence polarization measurements with TAMRA-labeled targets indicate 6.1 nM binding to PTH and only weak binding to off-target PTH related peptide (PTHrp). **(b)** Thread target sequence and redesign: threading peptides onto pseudorepetitive protein scaffolds. Left: schematic illustration. Right: Design model of SCT based on repeat protein scaffold (grey) and SCT target (orange). Fluorescence polarization measurements with TAMRA-labeled targets indicate 3.95 nM binding to SCT and 12 nM binding to GCG. **(c)** Binder design with deep network hallucination. Top left: schematic illustration. Right, designed binder resulting from Monte Carlo optimization of binder sequence using AlphaFold over 5000 steps, with only target sequence (not structure) provided. Hallucinated binder (gray); target Bid peptide (blue). Isothermal titration calorimetry measurements (far right) indicate 25 nM binding to Bid. Bottom: hallucination trajectory starting from random sequence (left) to final sequence (right); the protein folds around the peptide, which increases in helical content from step 0 to step 1000.

As an alternative to *de novo* parametric design of scaffolds that contain grooves, we explored the threading of helical peptides of interest onto already existing designed scaffolds with interfaces that make extensive interactions with helical peptides (Fig. 2b). We started from a library of scaffolds that contained single helices bound by pseudorepetitive helical scaffolds. We then threaded sequences of peptides of interest onto the bound single helix and filtered to maximize interfacial hydrophobic interactions of the target sequence to the binder scaffold. The binders were then redesigned in the presence of the threaded target sequence with ProteinMPNN^15^ and the complex was predicted with AF2^16^ (with *initial guess*^6^) and filtered on AF2 and Rosetta metrics.

Initial screening using yeast surface display identified 4/66 binders, which were expressed in *E. coli*. Following size exclusion chromatography (SEC) purification of the monomer fraction, all 4 of the designs were found to bind with sub-micromolar affinity using fluorescence polarization (FP), with the highest-affinity design binding with an affinity of 2.7 nM for SCT. Binding specificity was assessed with FP by measuring affinity for GCG, a related hormone to which SCT shares a significant degree of sequence identity (44%) and conformational homology^1,2^. We found that the tightest SCT binder was only 4 fold selective for SCT over GCG, which suggested additional design strategies might be necessary to increase the quality of the binding interface and to achieve high-specificity binding (Fig. 2b).

### Designing peptide binders by hallucination

We next explored the use of deep learning hallucination methods to generate helical peptide binders completely *de novo*, with no pre-specification of the desired binder geometry (from peptide sequence alone) (Fig. 2c). Hallucination or “activation maximization” approaches start from a network that predicts protein structure from sequence and carry out an optimization in sequence space for sequences which fold to structures with desired properties. This approach has been used to generate novel monomers^17^, functional-site scaffolds^14^ and cyclic oligomers^18^. Hallucination using AlphaFold2 (AF2) or RosettaFold has a number of attractive features for peptide binder design. First, neither the binder *nor the peptide* structure needs to be specified during the design process, enabling the design of binders to peptides in different conformations (this is useful given the unstructured nature of many peptides in solution; disordered peptides have been observed to bind in different conformations to different binding partners^9^). Second, metrics such as the predicted alignment error (pAE) have been demonstrated to correlate well with protein binding^6^, permitting the direct optimization of the desired objective, albeit with the possible hazard of generating adversarial examples^18^.

We began by designing binders to the apoptosis-related BH3 domain of Bid (Fig. 1a). The Bid peptide is unstructured in isolation, but adopts an alpha-helix upon binding to Bcl-2 family members^19,20^; it is therefore a model candidate for the design of helix-binding proteins. Starting from only the Bid primary sequence, and a random seed binder sequence (of lengths 60, 70, 80, 90 or 100 residues), we iteratively optimized the sequence of the binder through a Monte Carlo search in sequence space, guided by a composite loss function including the AF2 confidence (pLDDT, pTM) in the complex structure, and in the interaction between peptide and target (pAE). The trajectories typically converged in 5000 steps (sequence substitutions; Supplementary Fig. S3), and the output binder sequence was subsequently redesigned with ProteinMPNN, as previously described^18^. All designed binders were predicted to bind to Bid in a helical conformation; the exact conformations differ between designs because only the amino acid sequence of the target is specified in advance. This protocol effectively carries out flexible backbone protein design, which can be a challenge for traditional Rosetta based design approaches for which deep conformational sampling can be very compute intensive. Interestingly, in line with our prediction that “groove” scaffolds would offer an ideal topology for helical peptide binding, many of the binders from this approach contained a well-defined “groove” by eye, with the peptide predicted to make extensive interactions with the binder, typically helix-helix interactions.

47 of the hallucinated designs were tested experimentally (Supplementary Fig. S4a). Initial screening was performed with co-expression of a GFP-tagged Bid peptide and the HIS-tagged binders, with coelution of GFP and binder used as a readout for binding. 4 of these designs were further characterized, and showed soluble, monomeric expression even in the absence of peptide co-expression (Supplementary Fig. S4b). All designed proteins could be pulled-down using Bid BH3 peptide immobilized on beads (Supplementary Fig. S4c). Circular dichroism experiments indicated that the Bid peptide was unstructured in solution, and that helicity increased upon interaction with the hallucinated proteins, in line with the design prediction (Supplementary Fig. S4d). The binders were highly thermostable, and, unlike the native Bcl-2 protein Mcl-1, readily refolded after (partial) thermal denaturation at 95 °C (Supplementary Fig. S4e). Isothermal titration calorimetry revealed that all four bound Bid peptide, with the highest-affinity design binding having an affinity of 25 nM (Fig. 2c), a higher affinity interaction than with the native partner Mcl-1 (Supplementary Fig. S4f).

### Peptide binder design with RF*diffusion*

We next explored the design of binders using the RoseTTAFold-based denoising diffusion model RF*diffusion* described in the accompanying paper (Watson *et al*.). RF*diffusion* is much more compute efficient than hallucination, and is trained to directly generate a diversity of solutions to specific design challenges starting from random 3D distributions of residues that are progressively denoised. We reasoned that RF*diffusion* could be used both for binder optimization (by sampling related conformations around a specific binder structure) and for fully *de novo* design starting from a completely random noise distribution.

A long-standing challenge in protein design is to increase the activity of an input native protein or designed protein by exploring the space of plausible closely related conformations for those with predicted higher activity. This is difficult for traditional design methods as extensive full atom calculations are needed for each sample around a starting structure (using molecular dynamics simulation or Rosetta full atom relaxation methods), and it is not straightforward to optimize for higher binding affinity without detailed modeling of the binder-target sidechain interactions. We reasoned that, in contrast, RF*diffusion* might be able to rapidly generate plausible backbones in the vicinity of a target structure, increasing the extent and quality of interaction with the target guided by the extensive knowledge of protein structure inherent in RoseTTAfold. During the reverse diffusion (generative) process, RF*diffusion* takes random Gaussian noise as input, and iteratively refines this to a novel protein structure over many (“T”) steps (typically 200). Partly through this denoising process, the evolving structure no longer resembles “pure noise”, instead resembling a “noisy” version of the final structure. We reasoned that ensembles of structure with varying extents of deviation from an input structure could be generated by partially noising to different extents (for example, timestep 70), and then denoising to a similar, but not identical final structure (Fig. 3a, b).

**Figure 3.**
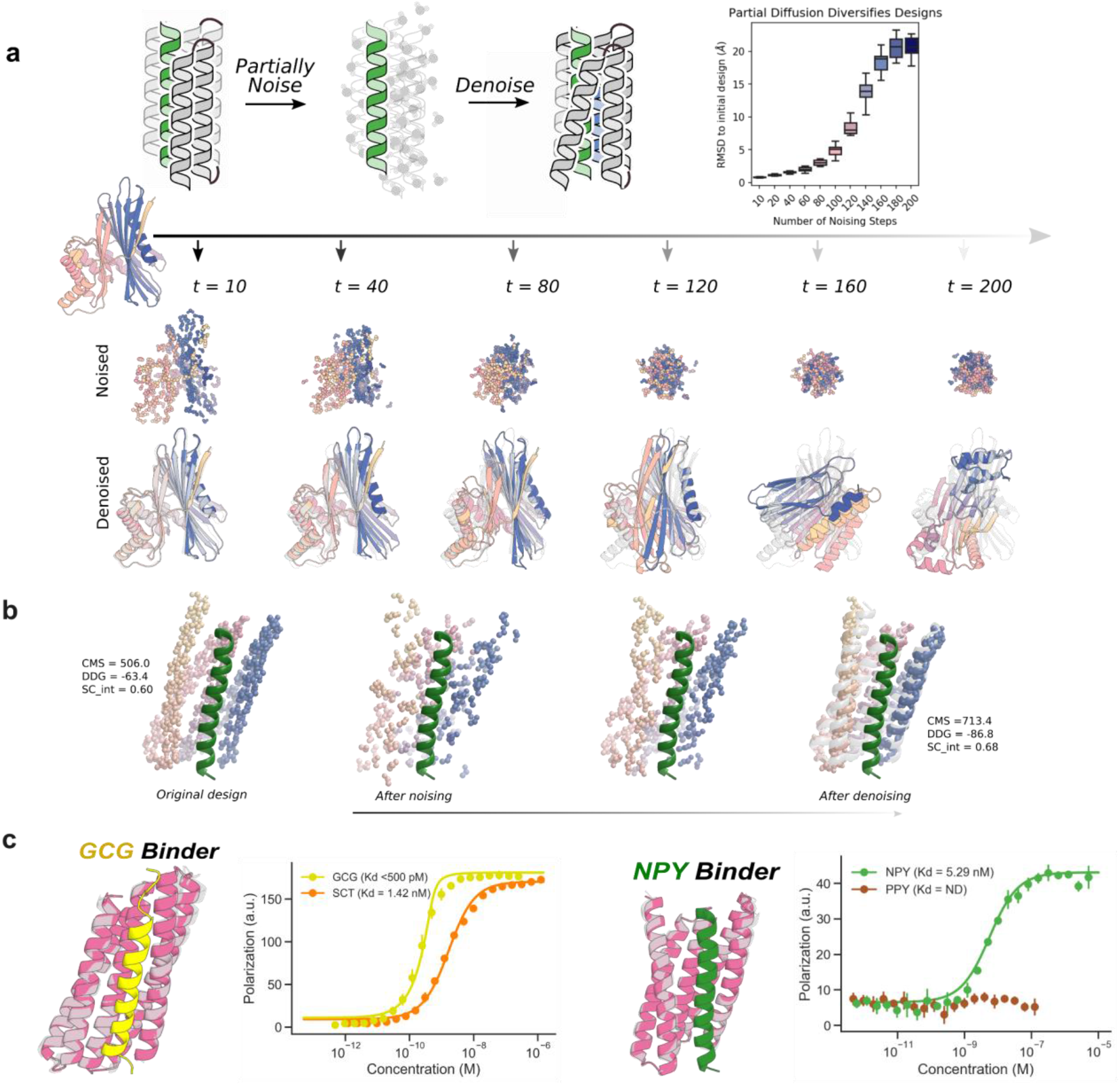
Peptide binder optimization with RF*diffusion*: **(a)** Top: Schematic showing partial noising and denoising using RF*diffusion*. A starting monomer (left) is partially noised for an increasing number of steps and then denoised resulting in designs (color) increasingly different from the original design (gray). Varying the noising stage from which denoising trajectories are initiated enables control over the extent of introduced structural variation. Bottom left: The distribution of RMSD to initial design vs number of partial noising steps. Bottom right: Starting from initial helix binder designs, we use partial diffusion to design optimized binders with improved shape complementarity. **(b)** Partial denoising trajectory starting from an initial NPY binder shown on the left. The final design (color) is shown on the right overlaid over the original design (gray). Contact molecular surface (CMS), Rosetta DDG (DDG) and interface shape complementarity (sc_int) values are reported for the original and optimized binder. **(c)** Diffused binders to GCG and NPY. Top left: Design models (gray) and AF2 predictions (pink, metrics in Supplementary Table 1), of diffused binders to GCG (yellow). Top right: FP measurements with FAM-labeled GCG indicate a sub-nanomolar binding affinity and selectivity over SCT. Bottom left: Design models (gray) and AF2 predictions (pink, metrics in Supplementary Table 1), of diffused binders to NPY (green). Bottom right: FP measurements with FAM-labeled NPY indicate a binding affinity of 5.29 nM and no binding to PYY, demonstrating selectivity.

We experimented with this approach starting from our parametrically-designed inpainted binders to GCG (with 231 nM affinity) and NPY (with 3.5 μM affinity) (Supplementary Fig. S2). Following partial noising and denoising, we identified designs that *in silico*, had significantly improved AF2 metrics compared to the starting design. The diversity compared to the starting design could be readily tuned by varying the time point to which the starting design was noised (Fig. 3a). Initial screening on yeast display revealed quite high binding success rates, with 25/96 designs binding GCG, and 20/96 binding NPY at 10 nM peptide concentration. The highest affinity designs were expressed in *E. coli*, purified, and their binding affinities were determined using FP. The highest-affinity binders were found to bind at subnanomolar affinities to GCG, and 5.6 nM to NPY (Fig. 3c). The designed proteins are quite specific: the GCG binders bound 10 times less tightly to SCT, which was chosen due to its high similarity to GCG. Impressively, the NPY binder did not show any cross-reactivity to peptide YY (PYY), which is a member of the NPY/pancreatic polypeptide family^21^ and shares a high percentage of sequence similarity (63.5% for the sequences used in the assay).

Inspired by this success at *optimizing* binders with RF*diffusion*, we next tested its ability to design binders to a different BH3 peptide, Bim and PTH completely *de novo* through unconditional binder design - providing RF*diffusion* only with the sequence and structures of the two peptides in helical conformations, and leaving the topology of the binding protein and the binding mode completely unspecified (Fig. 4a). From this minimal starting information, RF*diffusion* generated designs predicted by AF2 to fold and bind to the targets with high *in silico* success rates. A representative design trajectory is shown for PTH in Fig. 4b and Supplemental Video 1; starting from a random distribution of residues surrounding the PTH peptide in a helical conformation, in sequential denoising steps the residue shifts to surround the peptide and progressively organize itself into a folded structure which cradles the peptide along its entire surface.

**Figure 4.**
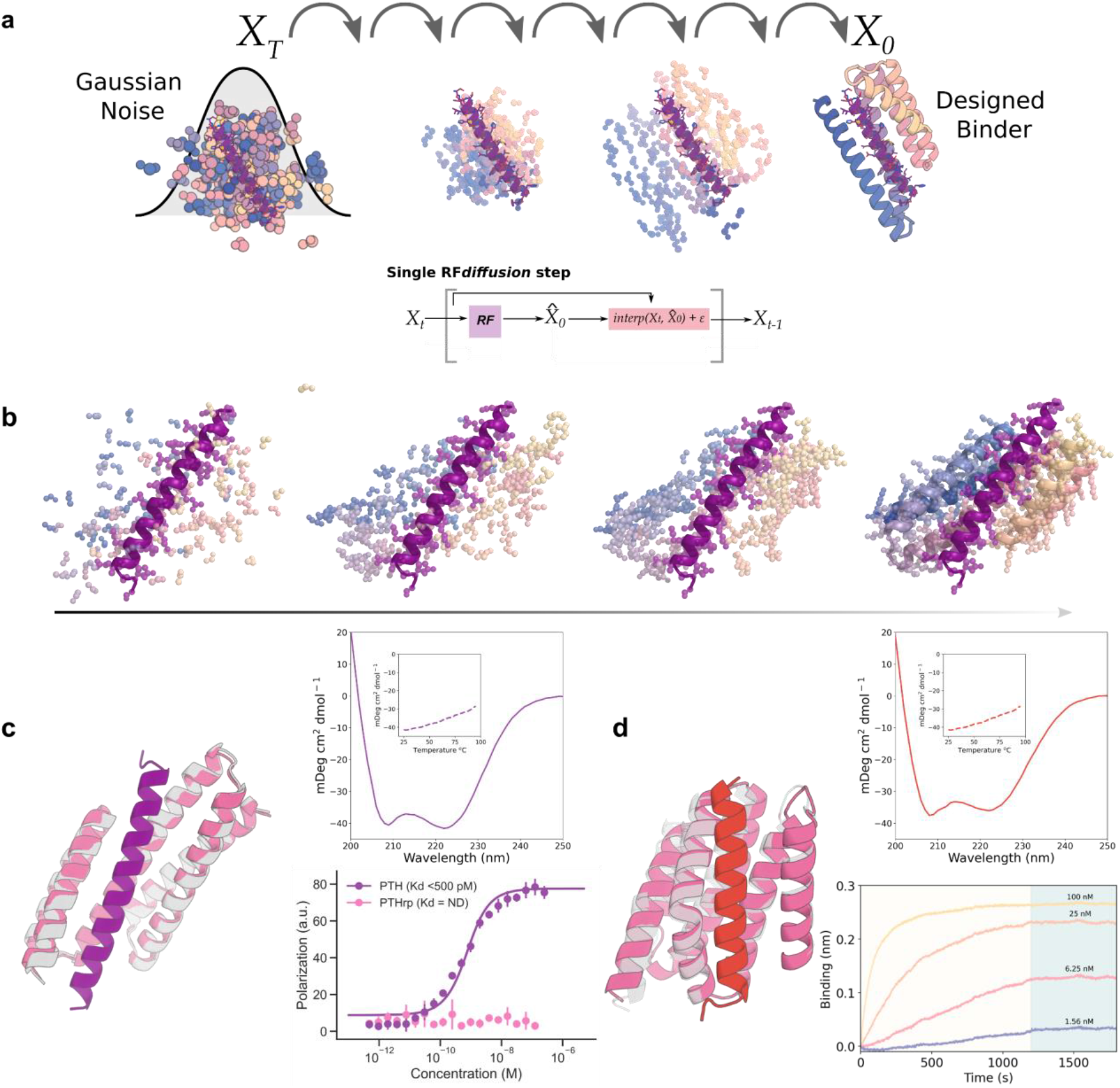
Peptide binder design with RF*diffusion*: **(a)** Schematic showing binder design using RFdiffusion. Starting from a random distribution of residues around the target peptide (X_T_), successive RFdiffusion denoising steps progressively remove the noise leading by the end of the trajectory X_0_ to a folded structure cradling the peptide. At each step t, RF*diffusion* predicts the final structure pX_0_ given the current noise sample Xt, and a step that interpolates in this direction is taken to generate the input for the next denoising step Xt-1. **(b)** Denoising trajectory in the presence of PTH (purple, Supplementary Video 1). Starting from random noise (left), a folded structure starts to emerge, leading to the final designed binder (right). **(c)** Design of picomolar affinity PTH binders. Left: Design model (gray) and AF2 prediction (pink, metrics in Supplementary Table 1), of designed PTH binder (purple). Bottom right: Fluorescence polarization measurements with TAMRA-labeled PTH indicate a sub-nanomolar binding affinity and no binding for PTH related peptide, indicating high specificity (PTHrp). Top right: Circular dichroism data indicating that the binder has the designed helical secondary structure and does not undergo cooperative unfolding below 95°C (inset). **(d)**: Design of picomolar affinity Bim binders. Left: Design model (gray) and AF2 prediction (pink, metrics in Supplementary Table 1), of designed Bim binder (red). Right bottom: Biolayer interferometry measurement of Bim binding indicates a sub-nanomolar affinity, with very slow dissociation kinetics. Biotinylated Bim was coupled to an Octet sensor, and incubated with the indicated concentrations of binder. The off rate is too slow to be accurately measured. Right top: CD data shows that the binder has helical secondary structure and is stable at 95°C (inset).

We obtained synthetic genes encoding 96 designs for each target. Using yeast surface display, we found that 25 of the 96 designs bound to Bim at 10 nM peptide concentration. The highest affinity design, which purified as a soluble monomer, bound too tightly for steady state estimates of the dissociation constant (Kd); global fitting of the association and dissociation kinetics suggest a Kd of ∼100pM (Fig 4C). For PTH, we found that 56/96 of the designs bound by yeast surface display with sub-micromolar affinities. The highest affinity design again bound too tightly for accurate Kd estimation; instead FP data provides an approximate upper bound for the Kd<500 pM (Fig. 4c). Binding was also highly specific; no binding was observed to the related PTHrp (Fig. 4c). Circular dichroism temperature melts indicate that both binders are stable at 95°C (Fig 4C). The diffused from scratch binders again had considerable structural similarity to our starting groove binding concept.

### Origins of higher affinity binding

The RF*diffusion* scaffolds bind the peptides with extended helices in a manner not entirely different from our starting groove structures and the other designs described above. What is the origin of their higher affinity? Reasoning that *de novo* building of the designs in the presence of the target, rather than starting from pre-generated scaffolds, could increase the extent of shape matching between binder and target, we computed the contact molecular surface^5^ for all of our designs in complex with the peptides. The average contact molecular surface for the partially diffused GCG binders and NPY increased by 33% and 29% respectively compared to the starting models, and the Rosetta ddG improved by 29% and 21% (Fig. S5a, S5b).

### Comparison of solutions to the binding problem

Our results provide an interesting side by side comparison of human and machine based problem solving. Despite the differences in affinity, the deep learning methods typically came up with the same overall solution to the helical peptide binding design problem–groove shaped scaffolds with helices lining the binding site–as the human designers did in the first Rosetta parametric approaches. The increased affinity likely derives at least in part from higher shape complementarity resulting from direct building of the scaffold to match the peptide shape; the ability of RF*diffusion* to “build to fit’’ provides a general route to creating high shape complementary binders to a wide range of target structures.

### Design of protein biosensors for PTH detection

Given our success in generating *de novo* binders to clinically-relevant helical peptides, we next sought to test their use as detection tools for use in diagnostic assays. Compared to immunosensors, which often exhibit antibody denaturation, loss of conformational stability, and wrong positioning of the antigen-binding site during sensor immobilization, *de novo* protein-based biosensors offer a more robust platform with high stability and tunability for diagnostics^22,23^. To design PTH biosensors, we grafted the 6.1 nM PTH binder into the lucCage system^24^, screened 8 designs for their luminescence response in the presence of PTH, and identified a sensitive lucCagePTH biosensor (LOD = 10 nM) with ∼21-fold luminescence activation in the presence of PTH (Fig. 5a).

**Figure 5.**
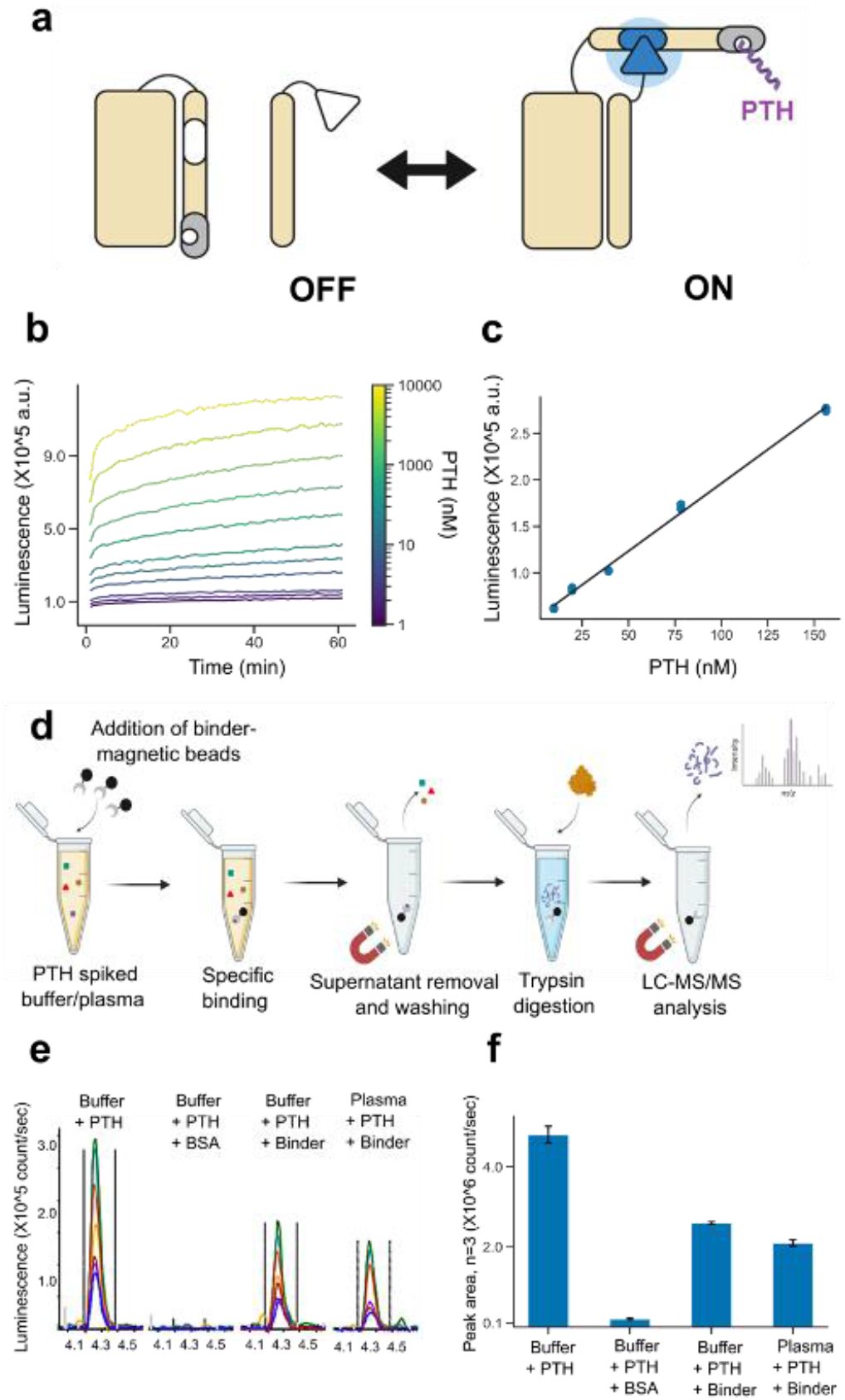
Application of designed binders to sensing and detection. **(a)** Protein biosensors for PTH detection. Left: Schematic of the grafted PTH lucCage biosensor, depicting the cage and latch (left, beige), key (right, beige), luciferase halves (inactive in white, active in blue), the PTH binder (red), and PTH peptide target (purple). Right: design model shown in the same color scheme. **(b)** Titration of PTH results in linear increases in luciferase luminescence. **(c)** Evaluation of the PTH biosensor at limiting concentrations of PTH indicates a 10 nM limit of detection (see methods). **(d-f)** The designed PTH binder enables robust recovery of PTH from complex mixtures. **(d)** Enrichment experiment schematic. **(e)** LC-MS/MS chromatograms for SVSEIQLMHNLGK, the N-terminal tryptic peptide of PTH; different peptide fragments detected by the LC-MS/MS assay are in different colors. **(f)** Mean chromatographic peak areas for triplicate measurements of each sample type. Error bars represent standard deviation.

### Enriching peptide targets from a complex mixture

We explored the use of our picomolar affinity *RFdiffusion* generated binder to PTH as a capture reagent in immunoaffinity enrichment coupled with liquid chromatography-tandem mass spectrometry (LC-MS/MS), a powerful platform for detecting low-abundance protein biomarkers in human serum^25^. We evaluated the RF*diffusion* binder in an LC-MS/MS assay for PTH in serum. PTH enrichment was quantified based on the analysis of the N-terminal peptide of a tryptic digestion of PTH in human plasma^26–28^. (see Supplemental Materials). We found that the designed binder enabled capture of PTH from spiked buffer and spiked human plasma with recoveries of 53% and 43%, respectively (Fig. 5b). The very high thermal stability of the designed binders (Fig. 4c,d) suggests that bioactive peptide capture reagents could have much longer shelf lives than antibodies, and be amenable to harsher washing conditions enabling re-use of binder conjugated beads.

## Discussion

Antibodies have served as the industry standard for affinity reagents for many years, but their use is often hampered by variable specificity and stability^29,30^. For binding helical peptides, the computationally designed helical scaffolds described in this paper have a number of structural and biochemical advantages. First, the extensive burial of the full length of an extended helix is difficult to accomplish with antibody loops, but very natural with matching extended alpha helices in groove shape scaffolds. Second, designed scaffolds are more amenable to incorporation into sensors as illustrated by the LucCage PTH sensor. Third, they are more stable, can be produced much less expensively, and could be more easily incorporated into affinity matrices for enrichment of peptide hormones from human serum. Fourth, peptide binders can achieve high affinity and specificity purely through computational methods, eliminating the need to use animals, which often mount weak responses to highly conserved bioactive molecules. Our MS based detection of peptides present at very low abundance in sera following enrichment using the designed binders could provide a general route forward for serological detection of a wide range of disease associated peptide biomarkers.

Our results highlight the emergence of powerful new deep learning methods for protein design. The inpainting and RF*diffusion* methods were both able to improve on initial Rosetta designs, and the hallucination approach generated high affinity binders without requiring prespecification of the bound structures. Most impressively, the RF*diffusion* method rapidly generated very high (picomolar) affinity and specific binders to multiple helical peptides. As described in the accompanying manuscript (Watson *et al*), RF*diffusion* is able to design binders to folded targets; here we demonstrate further that RF*diffusion* can be used to improve starting designs by partial noising and denoising, and can generate binders to peptides starting from no information other than the target. To our knowledge, the Bim and PTH binding proteins diffused starting from random noise are the highest affinity binders to any target (protein, peptide, or small molecule) achieved directly by computational design with no experimental optimization. We expect both the *de novo* peptide binder design capability and the ability to resample around initial designs (before or after experimental characterization) to be broadly applicable.

## Acknowledgements

This work was supported with funds provided by a grant U19 AG065156 from the National Institute for Aging (S.V.T., M.M., E.H., A.H., H.H.H., I.L., D.B.), a gift from Amgen (J.W.), the Audacious Project at the Institute for Protein Design (A.H.-W.Y., D.B.), a gift from Microsoft Gift supporting Computational Protein Structure Prediction and Design at the Institute for Protein Design (D.J., D.B.), the Washington State General Operating Fund supporting the Institute for Protein Design (P.V.), a grant INV-010680 from the Bill and Melinda Gates Foundation Grant (D.J., J.W., D.B.), a NIH NIBIB Pathway to Independence Award (A.H.-W.Y., K99EB031913), a National Science Foundation Training Grant number EF-2021552 (P.L.), NERSC award BER-ERCAP0022018 (P.L.), the Open Philanthropy Project Improving Protein Design Fund (P.L., G.R.L.,D.B.), The Donald and Jo Anne Petersen Endowment for Accelerating Advancements in Alzheimer’s Disease Research (N.Ben.), and the Howard Hughes Medical Institute (D.B.). J.M.R. and F.H. were supported by the Novo Nordisk Foundation (NNF19OC0054441 to J.M.R.). H.H.H is supported by a postdoctoral fellowship provided by the Partnership for Clean Competition. We thank Microsoft and AWS for generous gifts of cloud computing resources. We thank Ian Haydon for helping with figures and rendering. We thank Inna Goreshnik for supporting with chip assembly for yeast display experiments.

## Author Contributions

D.B. directed the work. I.L. and S.V.T. designed, screened, and experimentally characterized the parametrically designed groove scaffold peptide binders. P.J.Y.L. and S.V.T. designed, screened, and experimentally characterized the threaded peptide binders. J.L.W., developed the hallucination method for peptide binding. J.L.W., F.H., and J.M.R designed and experimentally characterized the hallucinated peptide binders. J.L.W. and S.V.T. designed and characterized the inpainted binders. S.V.T. and P.V. designed, screened, and experimentally characterized all the different classes of diffused peptide binders shown in this manuscript. J.L.W., D.J., and N.R.B. developed the RF*diffusion* algorithm used for peptide binder design. H.H.H, E.H., M.J.M., and A.N.H performed the LC-MS/MS peptide detection. A.H.-W.Y. designed and characterized the lucCagePTH biosensors and analyzed the sensing experiments. M.E. and G.R.L supported during yeast display binding screening. All authors reviewed and accepted the manuscript.

## Notes

https://www.bakerlab.org/wp-content/uploads/2022/11/diffusion_animation_PTHbinder.gif

### Competing Interest Statement

The authors have declared no competing interest.

### Summary of Updates

Increased figure resolution

