## Supplementary Methods for "De novo design of high-affinity protein binders to bioactive helical peptides"

### Gene construction of peptide hormone binders

The designed protein sequences were optimized to be both expressed in *S.cerevisiae* and *E. coli*. Linear DNA fragments (eBlocks, Integrated DNA Technologies) encoding design sequences included overhangs suitable for cloning into pETcon3 vector for yeast display[5] and Golden Gate cloning into LM627 vector for protein expression[18]. For initial testing hallucinated binders to Bid, binders were cloned into a modified LM627 vector. Specifically, Golden Gate cloning was used to generate sfGFP-Bid-STOP-[Binder]-SNAC-HISx6 assemblies.

### Yeast display screening

For the yeast transformation, 50-60 ng of digested pETcon3 and 100 ng of insert (eBlocks, Integrated DNA Technologies) were transformed into *S. cerevisiae* EBY100 strain using the protocol described in ref[5] . EBY100 cultures were grown in C-Trp-Ura medium supplemented with 2% (w/v) glucose (CTUG). For induction of expression, yeast cells initially grown in CTUG were transferred to SGCAA medium supplemented with 0.2% (w/v) glucose and induced at 30 °C for 16–24 h. Cells were washed with PBSF (PBS with 1% (w/v) BSA) and labeled for 40 minutes with biotinylated peptide targets at room temperature using without-avidity labeling condition[5]. After incubation time, cells were washed and resuspended in PBSF for cell sorting (Attune NxT Flow Cytometer, Thermo Fisher Scientific).

### NanoBiT screening

Linear gene fragments encoding binder design sequences and target peptide sequences were cloned into *E. coli* expression vectors using Golden Gate assembly; these vectors were pET28b(+) derivatives genetically fusing the smBiT and lgBiT halves of the NanoLuc® Luciferase (Promega) to the binders and peptides respectively. Resulting plasmids were transformed into BL21* (DE3) (Invitrogen) *E. coli* competent cells, then grown in 1mL TBII in 96-deepwell plates at 37C and 600 rpm. After 2 hours, expression was induced with IPTG (0.1 mM) and cells were incubated for an additional 4 hours. Cells were harvested by centrifugation (15 min at 4 kg), then resuspended in 100 uL lysis buffer (10 mM NaP pH 7.4, 150 mM NaCl, 5 mM $MgCl_2$, 1 mg/mL lysozyme, 10 ug/mL DNAse I, 1 tablet Complete Protease inhibitor / 50 mL). Cells were incubated for 1 hour at room temperature and 600 rpm, then frozen (-80C for 30min) and thawed (37C at 600 rpm for 30min) twice. Lysate was cleared by centrifugation (20 min at 4 kg), and the soluble fraction was then transferred to a 96-well plate for use as stock protein/peptide for conducting the nanoBiT screen. Screens were assembled in 96-well Half Area Black Flat Bottom Polystyrene NBS Microplates (Corning 3686). Binder design smBiT lysate was diluted 12 uL into 1400 uL assay buffer (10 mM NaP pH 7.4, 150 mM NaCl), while target peptide lgBiT lysate was diluted 6 uL into 1400 uL assay buffer. Stock rows in the assay plate were prepared by mixing 40 uL substrate (499.2 uL assay buffer, 20.8 uL Nano-Glo® Luciferase Assay Substrate (Promega)) with 40 uL diluted binder design smBiT lysate, while experimental rows were prepared by adding

50 uL diluted target peptide lgBiT lysate. At read time, 50 uL of the stock row was added to the 50 uL experimental row and mixed quickly and carefully, then luminescence was read immediately for 5 min using a plate reader (Biotek Synergy Neo2).

**Bicistronic protein expression**

Hallucinated binders to Bid were screened by bicistronic expression with the Bid peptide. Plasmids encoding sfGFP-Bid-STOP-[Binder]-SNAC-HISx6 were cloned into *E. coli,* and 2 mL cultures of each of the 47 designs were grown overnight in LB. Cultures were diluted into TB medium, and grown to approximately OD280 0.6, before induction with 1 mM IPTG for 4 hours at 37$^o$C. Bacteria were lysed for 15 minutes in 300 B-PER (Thermo) + 1 mM PMSF, 0.1 mg/mL Lysozyme (Sigma), 0.01 mg/mL DNAse I. Lysates were clarified by centrifugation at 4000 *g* for 10 minutes, before purification on Ni-NTA resin (wash buffer: 20 mM Tris pH 8.0, 150 mM NaCl, 20 mM Imidazole; elution buffer: 20 mM Tris pH 8.0, 150 mM NaCl, 250 mM Imidazole). Eluates were assessed for GFP fluorescence on a fluorescence plate reader.

**Peptide synthesis and purification**

The PTH-TAMRA peptide was synthesized in-house on a CEM Liberty Blue microwave synthesizer. All L- and D-amino acids were purchased from P3 Biosystems. Oxyma Pure was purchased from CEM, DIC was purchased from Oakwood Chemical, diisopropyl ethylamine (DIEA) and piperidine were purchased from Sigma- Aldrich. Dimethylformamide (DMF) was purchased from Fisher Scientific and treated with an Aldraamine trapping pack prior to use. Synthesis was done on a 0.1 mmol scale on CEM Cl-TCP(Cl) resin. Five equivalents of each amino acid were activated using 0.1 M Oxyma with 2% (v/v) DIEA in DMF, 15.4% (v/v) DIC, and coupled on resin for 4 min with double coupling if needed. This was followed by deprotection using 5 mL of 20% piperidine in DMF for 2 min at 95 °C. Global deprotection was accomplished TFA/Water/TIPS (95:2.5:2.5) for 3 hours. This deprotection mixture was precipitated in 30 mL of ice-cold ethyl ether, centrifuged and decanted, then washed twice more with fresh ether and dried under nitrogen to yield crude peptide for high pressure liquid chromatography (HPLC) purification.

The crude peptide was dried and dissolved in a mixture of ACN and water where the entire crude is soluble. This solution was purified on a C18 column in an Agilent HPLC instrument. A linear gradient of increasing ACN with 0.1% TFA was used to purify the samples. UV signal was monitored at 214 nm and all peaks were collected. Peaks were checked using ESI mass spectroscopy for the correct peptide mass. The purified peptide was then lyophilized for further use.

**Protein expression and purification in *E. coli* for peptide hormone binders**

Protein expression was performed using 50 mL of the Studier autoinduction media supplemented with kanamycin, and grown overnight at 37°C. The cells were harvested by spinning at 4,000 x g for 10 min and then resuspended in lysis buffer (100 mM Tris-HCl, 200 mM NaCl, 50 mM imidazole) supplemented with protease inhibitor tablets (Pierce™ Protease Inhibitor Tablets, EDTA-free). Then, the cells were lysed by sonication in a Qsonica, Q500 with a: 4-pronged horn for 2:30 min ON total, with an amplitude of 80%. Soluble fractions were clarified by centrifugation at 14,000 x g for 40 minutes, and were subsequently purified by affinity chromatography using bed Ni-NTA resin (Qiagen or Thermo Fisher) on a vacuum manifold. A series of washes using Low-salt buffer (20 mM Tris-HCl, 200 mM NaCl, 50 mM imidazole) and High- salt buffer (20 mM Tris-HCl, 1000 mM NaCl, 50 mM imidazole) were performed prior to elution with Elution buffer (20 mM Tris-HCl, 200 mM NaCl, 500 mM imidazole). After elution, protein samples were filtered and injected into an autosampler-equipped Akta pure system on a Superdex S75 Increase 10/300 GL column at room temperature. The SEC running buffer was 20mM Tris-HCl, 100mM NaCl pH 8. Selected fractions were pooled and concentrated using Spin filters (3 kDa molecular weight cutoff, Amicon, Millipore Sigma) and stored at 4 °C before downstream characterizations. Protein concentrations were determined by absorbance at 280 nm using a NanoDrop spectrophotometer (Thermo Scientific) using their extinction coefficients and molecular weights obtained from their amino acid sequences using the ProtParam tool.

**Fluorescence polarization**

Fluorescence polarization binding assays were carried out in 96-well plates (Corning 3686), with two-fold serial dilution of designed peptide binders in the presence of 0.5 nM fluorescently labeled peptide targets. Protein and peptide were diluted from their stock concentration into 20mM Tris-HCl pH 8, 100mM NaCl, 0.1% v/v Tween 20, and the protein was titrated in 2-fold serial dilutions onto constant peptide. After incubating the peptide and binder for one hour at room temperature, the fluorescence polarization was measured at the excitation and emission wavelengths of the FAM dye (485/530 nm) or the TAMRA dye (530/590 nm), in a Synergy Neo2 multi-mode plate reader. Titrations were conducted in replicate, and the Kd was fitted with SciPy. Specifically, curves were fit to $N$ observations of an observed signal, $Signal_i$, at titrated concentrations $[A_{tot}]_i$ according to the following equation:

$$Signal_i = Baseline + Amplitude \frac{AB_{conc}([A_{tot}]_i, [B_{tot}], K_d)}{[B_{tot}]},$$

Where $[B_{tot}]$ is the known total concentration of the binder, $Baseline$ and $Amplitude$ are free parameters, and the concentration of the bound state $[AB]$ is computed as

$$AB_{conc}([A_{tot}]_i, [B_{tot}], K_d) = (([A_{tot}] + [B_{tot}] + K_d) \pm \sqrt{([A_{tot}] + [B_{tot}] + K_d)^2 - 4[A_{tot}][B_{tot}]})/2$$

The unknown parameters ( $K_D$, $Baseline$ and $Amplitude$) were fit using scipy.optimize.curve_fit, $[B_{tot}]$was additionally fit in the optimization, but only allowed to within 0.5 nM $\pm 0.1\%$.

Peptides used for the assay are shown in Table 1.

**Table 1. Fluorophore-labeled peptides used in Fluorescence polarization assays**

| Peptide name | Sequence | Supplier | Cat # | Fluorophore |
|---|---|---|---|---|
| PTH-TAMRA | SVSEIQLMHNLGKHLNSMERVE WLRKKLQDVHNF | In-house | NA | 5-TAMRA |
| PTHrp-FAM | AVSEHQLLHDKGKSIQDLRRRF FLHHLIAEIHTAEIA | Phoenix Pharmaceu ticals, Inc. | FG-056-08A | FAM |
| SCT-FAM | HSDGTFTSELSRLREGARLQRL LQGLV | Phoenix Pharmaceu ticals, Inc. | FG-067-03A | FAM |
| GCG-FAM | HSQGTFTSDYSKYLDSRRAQD FVQWLMNT | Addex Bio | ABBFO 2033 | FAM |
| NPY-FAM | SKPDNPGEDAPAEDMARYYSA LRHYINLITRQR | Phoenix Pharmaceu ticals, Inc. | FG-049-04A | FAM |
| PPY-FAM | IKPEAAGEDASPEELNRYYASL RHYLNLVTRQRY | Phoenix Pharmaceu ticals, Inc. | FG-059-02A | FAM |

**Cloning, expression and purification of Bid-binding hallucinations, Avi-tagged Bid peptide and MCL-1**

Bid-binding hallucinations were cloned into a pET28 vector, containing an N-terminal $His_{10}$ and a PreScission cleavage site, using TEDA cloning[32] and transformed into XL-1-Blue chemically competent cells, single clones isolated and amplified and sequences confirmed by Sanger sequencing. Plasmids transformed into chemically competent BL21 DE3 *E. coli*, and plated onto LB agar plates supplemented with 100 ug/mL kanamycin. Single colonies were used to make starter cultures of LB with 100 ug/mL kanamycin and incubated overnight at 37 °C. 1:100 volume starter culture was added to autoinduction media Overnight Express Instant TB Medium (Novagen) in Ultra-Yield flasks (Thomson), with 100 µg/mL kanamycin, incubated at 37 °C for 5 hours, then 18 °C for 18 hrs. Cells harvested by centrifugation 6,000 rpm, 20 mins, 4 °C, and pellets were frozen at -80°C.

Defrosted cell pellets were resuspended in approx. 10 mL/g Lysis Buffer (50 mM potassium phosphate pH 7.0, 300 mM NaCl, 5 mM imidazole, 2 mM b-mercaptoethanol, 10% glycerol), supplemented with 60 µg/mL lysozyme, 1.4 µg/mL DNaseI, 0.05 mM PMSF. Cells were lysed by passing through French press twice, 18 kpsi. Lysate was clarified by centrifugation 18,000 *g*, 45 mins, 4 °C, and loaded onto HIS-Select Nickel affinity resin (Sigma) by gravity, resin washed with Wasg Buffer (50 mM potassium phosphate pH 7.0, 100 mM NaCl, 5 mM imidazole, 2 mM b-mercaptoethanol, 10% glycerol) and eluted with Wash Buffer containing 350 mM imidazole. Protein containing fractions (assessed by $A_{280}$) were combined, and further purified by size exclusion chromatography (SEC) using HiLoad 16/600 200 pg Superdex column (Cytiva) using ÅKTA FPLC system (Cytiva) equilibrated in 50 mM sodium phosphate pH 7.0, 1 mM DTT. Fractions were concentrated, concentration measured using $A_{280}$ and predicted extinction coefficients[33], then flash frozen $N_{2(l)}$ for storage at -80 °C.

DNA corresponding to BH3 motif of human Bid Q79-G144 (Uniprot: P55957) was assembled by complementary oligos (IDT) and primer extension using Klenow fragment (NEB), and cloned using TEDA into pET28 with an N-terminal $His_{10}$, SUMO and C-terminal Avi. Expression and purification was carried out as for the hallucinations, except for co-transformation with a chloramphenicol-resistant BirA expressing plasmid, the addition of chloramphenicol 25 ug/mL in all cultures, with the addition of 40 µM BTN to the media before temperature was reduced to 18 °C. After SEC, $His_{10}$-SUMO was cleaved using ULP-1 protease, and $His_{10}$-SUMO removed using Ni resin, Bid-Avi peptide concentration was measured using $A_{280}$, and stored at -80°C. To express human Mcl-1 P166-G327 (Uniprot: Q07820) a pEQ80L vector with N-terminal $His_6$ and Avi-tag, for co-expression with BirA. Expression and purification was carried out as for the Bid-binding hallucinations, with the addition of 40 µM BTN to the media before temperature was reduced to 18 °C.

**ITC**

Isothermal titration calorimetry was carried out with an ITC200 (Micocal). Bid peptide was in the syringe, at ~300 µM, and binder (hallucination of Mcl-1) was kept in the cell (~25 µM), with both peptide and binder in matched buffer (sodium phosphate pH 7.0, 1 mM DTT). Temperature was held at 25 °C or 10 °C, as indicated. Fitting of titrations was carried out using 1-site binding, using manufacturers software (OriginLab).

### Circular dichroism

Spectra were recorded for Bid peptide alone, Bid in complex with binders (hallucination or Mcl-1) and binders alone. All concentrations were 10 µM, in a 2 cm pathlength quartz cuvette. Spectra recorded on J-1500 Circular Dichroism Spectrophotometer, with temperature held at 25 °C, or ramped at 1 °C/min.

### Pull-down

10 µL bead slurry Dynabeads M-280 Streptavidin (Thermo Fisher Scientific) were washed with Pull-Down Buffer (sodium phosphate pH 7.0, 1 mM DTT, 0.05% Tween20), incubated with saturating amounts of (Avi-tagged) Bid peptide 15 mins, 4 °C with rotation, beads were then incubated with free biotin 25 µM, and washed three times with ice cold Pull-Down Buffer. 10 µL of 2 µM binder (hallucination or Mcl-1) was incubated with pelleted beads for 30 mins, 4 °C, with rotation. Supernatant was recovered and the beads washed three times before resuspension in 10 µL Pull-Down Buffer. Both supernatant and washed beads were loaded onto denaturing SDS-PAGE, with protein detection by InstantBlue Coomassie staining.

### Bio-layer Inferometry (BLI) Binding Experiments

BLI experiments were performed on an Octet Red96 (ForteBio) instrument, with streptavidin coated tips (Sartorius Item no. 18-5019). Buffer comprised 1X HBS-EP+ buffer (Cytiva BR100669) supplemented with 0.1% w/v bovine serum albumin. Tips were pre-incubated in the buffer for at least 10 minutes before use. Tips were then sequentially incubated in 50nM biotinylated Bim peptide (loading, 500s), buffer (baseline, 150s), designed binder (association, 1200s) and buffer (dissociation, 600s). Due to the extremely slow dissociation of Bim from the designed binders, it was not possible to calculate a precise $K_D$, but estimates suggest significantly sub-nanomolar affinity.

### Design and characterization of lucCagePTH biosensor for parathyroid hormone detection

The detailed design protocol for the lucCage and lucKey sensor system was described previously (Nature, 2021, 482). In brief, the amino acid sequence (FELLDKLIELLRELIELTREYI) at the N-terminal end of the 6.1 nM PTH binder was grafted onto the latch region (residues 323 to 353) of lucCage. The Rosetta models were visually inspected and eight of them were selected for experimental validation. We produced, purified, and screened for the luminescence signal emitted from each biosensor in the presence of 5 µM PTH. From this process, we identified several hits showing increased luminescence upon adding PTH, of which we assigned the best one with a 21-fold activation as lucCagePTH. We then set up assays to evaluate the response of lucCagePTH with a range of PTH concentrations. 10 µl of 10 nM lucCagePTH, 10 µl of 10 nM lucKey, 10 µl of serial diluted PTH, and 40 µl of buffer (50% HBS-EP/50% Nano-Glo luciferase assay buffer) were pre-mixed and 30 µl of 100× diluted furimazine was injected immediately before luminescence kinetic acquisition. The luminescence measurements were taken every 1 min (0.1 s integration and 10 s shaking during intervals) for a total of 60 mins by Neo2 microplate reader. The linear region of luminescence responses to the corresponding PTH concentrations was fitted to a linear regression curve and the LOD was calculated as 3 × standard deviation of the response / the slope of the calibration curve.

**Affinity enrichment of PTH analyzed by LC-MS/MS**

- **Sample description**

Recombinant human PTH protein was purchased from Sigma (#SAE 0192_100 ug, MA, USA) and reconstituted at 100 µg/mL in a 10 % acetonitrile, 0.1 % formic acid,1 mg/mL bovine serum albumin solution and stored in 40 µL aliquots at -20 °C. Dilutions at 1000 ng/mL and 62.5 ng/mL were prepared freshly as needed by dilution in the same acetonitrile, formic acid, albumin solution.

The plasma samples used were de-identified leftover clinical samples obtained from the clinical laboratories at the University of Washington Medical Center. The use of de-identified leftover clinical samples was reviewed by the University of Washington Human Subjects Division (STUDY00013706).

The evaluation of PTH immunoaffinity enrichment in buffer and plasma was performed in three process replicates using 8 different types of samples:

- Series A: Reconstitution buffer (10 % acetonitrile, 0.1 % formic acid,1 mg/mL bovine serum albumin in water) served as the blank.
- Series B: Reconstitution buffer spiked with PTH at 7.2 ng/mL was directly digested without the addition of beads and served as the Control sample (representing 100% recovery of PTH).
- Series C: Reconstitution buffer spiked with PTH at 7.2 ng/mL was incubated with beads blocked by bovine serum albumin before washing and digestion, which served as the negative control, to quantify non-specific binding in buffer.
- Series D: Reconstitution buffer spiked with PTH at 7.2 ng/mL was incubated with designed binder-conjugated beads before washing and digestion, which was used to quantify the affinity precipitation of PTH from buffer.
- Series E: Plasma was incubated with beads blocked by bovine serum albumin before washing and digestion, which was used to quantify non-specific binding in unspiked plasma.
- Series F: Plasma was incubated with designed binder-conjugated beads before washing and digestion, which was used to quantify affinity precipitation of PTH in plasma.
- Series G: Plasma spiked with PTH at 7.2 ng/mL was incubated with beads blocked by bovine serum albumin before washing and digestion, which was used to quantify non-specific binding in spiked plasma.
- Series H: Plasma spiked with PTH at 7.2 ng/mL was incubated with designed binder-conjugated beads before washing and digestion, which was used to quantify the affinity precipitation of PTH in spiked plasma.

- **Sample preparation and LC-MS/MS conditions**

Affinity enrichment was performed in buffer or plasma at the protein level. Designed binders were conjugated to tosyl-activated Dynabeads M-280 according to the manufacturer's instructions and subsequently blocked using bovine serum albumin and Tris. The amino terminal peptide was analyzed after tryptic digestion of either pure protein in buffer, or after trypsin digestion of PTH that had been affinity precipitated by the designed binder-conjugated beads (or by the control/blocked magnetic beads). Briefly, PTH proteins in buffer/plasma were purified using PTH mini-binder conjugated-paramagnetic beads at room temperature, for 1 h. The beads were then washed 4 times with phosphate-buffered saline supplemented with CHAPS (0.1% 3-((3cholomidopropyl) dimethylammonio)-1-propanesulfate to reduce nonspecific interactions). The proteins that were affinity precipitated by the designed binder-conjugated-paramagnetic beads were suspended in 10 µL of a solution containing 10 % acetonitrile, 0.1 % formic acid, 1 mg/mL bovine serum albumin. The washed beads were then suspended with 30 µL of 30% isopropanol,100 mM ammonium bicarbonate, and digested at 37 °C for 30 min after adding 100 µL of 0.01 mg/mL trypsin in 10 mM hydrochloride acid. The liberated peptides were then removed from the beads using a magnet and analyzed using LC-MS/MS.

Peptides were analyzed by liquid chromatography-tandem mass spectrometry in the multiple reaction monitoring acquisition mode using an UHPLC I-Class Chromatography system coupled to a Xevo TQ-S triple quadrupole tandem mass spectrometer (Waters, MA, USA). Peptides were eluted from an Acquity UPLC HSS T3 1.8µm (C18, 2.1x50 mm, pore size 100 Å) analytical column (Waters) at 45 °C using 0.1 % formic acid, 2 % dimethylsulfoxide in LC-MS grade water as mobile phase A and 0.1 % formic acid, 2 % dimethylsulfoxide in LC-MS grade methanol as mobile phase B.

The liquid chromatography and mass spectrometry conditions are detailed in Table 2, 3 and 4.

| **Table 2. Liquid chromatography conditions** | |
|---|---|
| Mobile phase | Phase A: 0.1 % formic acid, 2 % dimethylsulfoxide in water<br>0.1 % formic acid, 2 % dimethylsulfoxide in methanol |
| Column | Acquity UPLC HSS T3 1.8µm (C18, 2.1x50 mm, pore size 100 Å) |
| Temperature | 45±5 °C |
| Flow rate | 0.3 mL/min |
| Injection volume | 20 µL |
| Gradient | 0-0.5 min: 2% B at 0.3 mL/min<br>7.5: 98% B at 0.3 mL/min<br>7.6: 98% B at 0.6 mL/min<br>8.6: 2% B at 0.6 mL/min<br>9.9: 2% at 0.3 mL/min |

| Table 3. Mass spectrometry conditions | |
|---|---|
| Source polarity | ESI+ |
| Capillary voltage | 3.25 kV |
| Source Offset voltage | 50 V |
| Desolvation Temp | 600 °C |
| Desolvation Gas Flow | 1000 L/h |
| Cone Gas Flow | 150 L/h |

**Table 4. Multiple reaction monitoring conditions**

| Peptide sequences | Q1 (m/z) | Q3 (m/z) | Cone (V) | Collision Energy (eV) | Ion type |
|---|---|---|---|---|---|
| HLNSMER.2 | 443.7136 | 218.1047 | 35 | 15 | y3 |
| HLNSMER.2 | 443.7136 | 261.6207 | 35 | 15 | y4 |
| HLNSMER.2 | 443.7136 | 318.6421 | 35 | 15 | y5 |
| HLNSMER.3 | 296.1448 | 218.1047 | 35 | 9 | y3 |

| | | | | | |
|---|---|---|---|---|---|
| HLNSMER.3 | 296.1448 | 261.6207 | 35 | 9 | y4 |
| HLNSMER.3 | 296.1448 | 318.6421 | 35 | 9 | y5 |
| HLNSMER.3 | 296.1448 | 435.202 | 35 | 9 | y3 |
| HLNSMER.3 | 296.1448 | 522.2341 | 35 | 9 | y4 |
| HLNSMER.3 | 296.1448 | 636.277 | 35 | 9 | y5 |
| HLNSM(+15.994915)ER.2 | 451.7111 | 226.1021 | 35 | 16 | y3 |
| HLNSM(+15.994915)ER.2 | 451.7111 | 269.6181 | 35 | 16 | y4 |
| HLNSM(+15.994915)ER.2 | 451.7111 | 326.6396 | 35 | 16 | y5 |
| HLNSM(+15.994915)ER.2 | 451.7111 | 451.1969 | 35 | 16 | y3 |
| HLNSM(+15.994915)ER.2 | 451.7111 | 538.229 | 35 | 16 | y4 |
| HLNSM(+15.994915)ER.2 | 451.7111 | 652.2719 | 35 | 16 | y5 |
| HLNSM(+15.994915)ER.3 | 301.4765 | 226.1021 | 35 | 10 | y3 |
| HLNSM(+15.994915)ER.3 | 301.4765 | 269.6181 | 35 | 10 | y4 |

| | | | | | |
|---|---|---|---|---|---|
| HLNSM(+15.994915)ER.3 | 301.4765 | 326.6396 | 35 | 10 | y5 |
| HLNSM(+15.994915)ER.3 | 301.4765 | 451.1969 | 35 | 10 | y3 |
| HLNSM(+15.994915)ER.3 | 301.4765 | 538.229 | 35 | 10 | y4 |
| ADVNVLTK.2 | 430.2478 | 574.3559 | 35 | 15 | y5 |
| ADVNVLTK.2 | 430.2478 | 673.4243 | 35 | 15 | y6 |
| ADVNVLTK.3 | 287.1676 | 181.1259 | 35 | 9 | y3 |
| ADVNVLTK.3 | 287.1676 | 230.6601 | 35 | 9 | y4 |
| ADVNVLTK.3 | 287.1676 | 361.2445 | 35 | 9 | y3 |
| ADVNVLTK.3 | 287.1676 | 460.313 | 35 | 9 | y4 |
| SLGEADK.2 | 360.1821 | 167.0921 | 35 | 12 | y3 |
| SLGEADK.2 | 360.1821 | 231.6134 | 35 | 12 | y4 |
| SLGEADK.2 | 360.1821 | 260.1241 | 35 | 12 | y5 |
| SLGEADK.2 | 360.1821 | 333.1769 | 35 | 12 | y3 |

| | | | | | |
|---|---|---|---|---|---|
| SLGEADK.2 | 360.1821 | 462.2195 | 35 | 12 | y4 |
| SLGEADK.2 | 360.1821 | 519.2409 | 35 | 12 | y5 |
| SLGEADK.3 | 240.4572 | 167.0921 | 35 | 7 | y3 |
| SLGEADK.3 | 240.4572 | 260.1241 | 35 | 7 | y5 |
| SLGEADK.3 | 240.4572 | 333.1769 | 35 | 7 | y3 |
| SLGEADK.3 | 240.4572 | 462.2195 | 35 | 7 | y4 |
| VEWLR.2 | 351.7003 | 229.1183 | 35 | 12 | b2 |
| VEWLR.2 | 351.7003 | 474.2823 | 35 | 12 | y3 |
| EDNVLVESHEK.2 | 649.8148 | 629.2889 | 35 | 23 | y5 |
| EDNVLVESHEK.2 | 649.8148 | 728.3573 | 35 | 23 | y6 |
| EDNVLVESHEK.2 | 649.8148 | 841.4414 | 35 | 23 | y7 |
| EDNVLVESHEK.3 | 433.5456 | 315.1481 | 35 | 14 | y5 |
| EDNVLVESHEK.3 | 433.5456 | 364.6823 | 35 | 14 | y6 |

| | | | | | |
|---|---|---|---|---|---|
| EDNVLVESHEK.3 | 433.5456 | 421.2243 | 35 | 14 | y7 |
| EDNVLVESHEK.3 | 433.5456 | 629.2889 | 35 | 14 | y5 |
| DAGSQRPR.2 | 443.7281 | 322.1854 | 35 | 15 | y5 |
| DAGSQRPR.2 | 443.7281 | 643.3634 | 35 | 15 | y5 |
| DAGSQRPR.2 | 443.7281 | 700.3849 | 35 | 15 | y6 |
| DAGSQRPR.3 | 296.1545 | 214.6401 | 35 | 9 | y3 |
| DAGSQRPR.3 | 296.1545 | 278.6693 | 35 | 9 | y4 |
| DAGSQRPR.3 | 296.1545 | 322.1854 | 35 | 9 | y5 |
| DAGSQRPR.3 | 296.1545 | 350.6961 | 35 | 9 | y6 |
| DAGSQRPR.3 | 296.1545 | 428.2728 | 35 | 9 | y3 |
| DAGSQRPR.3 | 296.1545 | 556.3314 | 35 | 9 | y4 |
| DAGSQRPR.3 | 296.1545 | 643.3634 | 35 | 9 | y5 |
| DAGSQRPR.3 | 296.1545 | 700.3849 | 35 | 9 | y6 |

| | | | | | |
|---|---|---|---|---|---|
| SVSEIQLMHNLGK.2 | 728.3849 | 527.2973 | 35 | 26 | y9 |
| SVSEIQLMHNLGK.2 | 728.3849 | 568.3202 | 35 | 26 | y5 |
| SVSEIQLMHNLGK.2 | 728.3849 | 635.3346 | 35 | 26 | y11 |
| SVSEIQLMHNLGK.2 | 728.3849 | 699.3607 | 35 | 26 | y6 |
| SVSEIQLMHNLGK.2 | 728.3849 | 812.4447 | 35 | 26 | y7 |
| SVSEIQLMHNLGK.2 | 728.3849 | 940.5033 | 35 | 26 | y8 |
| SVSEIQLMHNLGK.2 | 728.3849 | 1053.587 | 35 | 26 | y9 |
| SVSEIQLMHNLGK.2 | 728.3849 | 1269.662 | 35 | 26 | y11 |
| SVSEIQLMHNLGK.3 | 485.9257 | 159.1128 | 35 | 16 | y3 |
| SVSEIQLMHNLGK.3 | 485.9257 | 431.2613 | 35 | 16 | y4 |
| SVSEIQLMHNLGK.3 | 485.9257 | 470.7553 | 35 | 16 | y8 |
| SVSEIQLMHNLGK.3 | 485.9257 | 527.2973 | 35 | 16 | y9 |
| SVSEIQLMHNLGK.3 | 485.9257 | 568.3202 | 35 | 16 | y5 |

| | | | | | |
|---|---|---|---|---|---|
| SVSEIQLMHNLGK.3 | 485.9257 | 591.8186 | 35 | 16 | y10 |
| SVSEIQLMHNLGK.3 | 485.9257 | 635.3346 | 35 | 16 | y11 |
| SVSEIQLMHNLGK.3 | 485.9257 | 699.3607 | 35 | 16 | y6 |
| SVSEIQLMHNLGK.3 | 485.9257 | 812.4447 | 35 | 16 | y7 |
| SVSEIQLMHNLGK.3 | 485.9257 | 940.5033 | 35 | 16 | y8 |
| SVSEIQLM(+15.994915)HNLGK.2 | 736.3823 | 159.1128 | 35 | 26 | y3 |
| SVSEIQLM(+15.994915)HNLGK.2 | 736.3823 | 317.2183 | 35 | 26 | y3 |
| SVSEIQLM(+15.994915)HNLGK.2 | 736.3823 | 431.2613 | 35 | 26 | y4 |
| SVSEIQLM(+15.994915)HNLGK.2 | 736.3823 | 535.2948 | 35 | 26 | y9 |
| SVSEIQLM(+15.994915)HNLGK.2 | 736.3823 | 568.3202 | 35 | 26 | y5 |
| SVSEIQLM(+15.994915)HNLGK.2 | 736.3823 | 643.3321 | 35 | 26 | y11 |
| SVSEIQLM(+15.994915)HNLGK.2 | 736.3823 | 715.3556 | 35 | 26 | y6 |
| SVSEIQLM(+15.994915)HNLGK.2 | 736.3823 | 828.4396 | 35 | 26 | y7 |

| | | | | | |
|---|---|---|---|---|---|
| SVSEIQLM(+15.994915)HNLGK.2 | 736.3823 | 956.4982 | 35 | 26 | y8 |
| SVSEIQLM(+15.994915)HNLGK.2 | 736.3823 | 1069.582 | 35 | 26 | y9 |
| SVSEIQLM(+15.994915)HNLGK.3 | 491.2573 | 159.1128 | 35 | 16 | y3 |
| SVSEIQLM(+15.994915)HNLGK.3 | 491.2573 | 643.3321 | 35 | 16 | y11 |

- **Data treatment**

Data processing was performed with Skyline Daily version 21.1.1.223. The peak area for each peptide was determined as the sum of the peak areas of all selected transitions. The recovery over blocked-beads (RE) in spiked buffer and in spiked plasma was estimated using Equations 1, and 2, respectively.

$$RE_{buffer} = \frac{Peak\ area\ Series\ D}{Peak\ area\ Series\ B} \quad (1)$$

$$RE_{plasma} = \frac{Peak\ area\ Series\ H}{Peak\ area\ Series\ B} \quad (2)$$
