## Supplementary Materials for "De novo design of high-affinity protein binders to bioactive helical peptides"

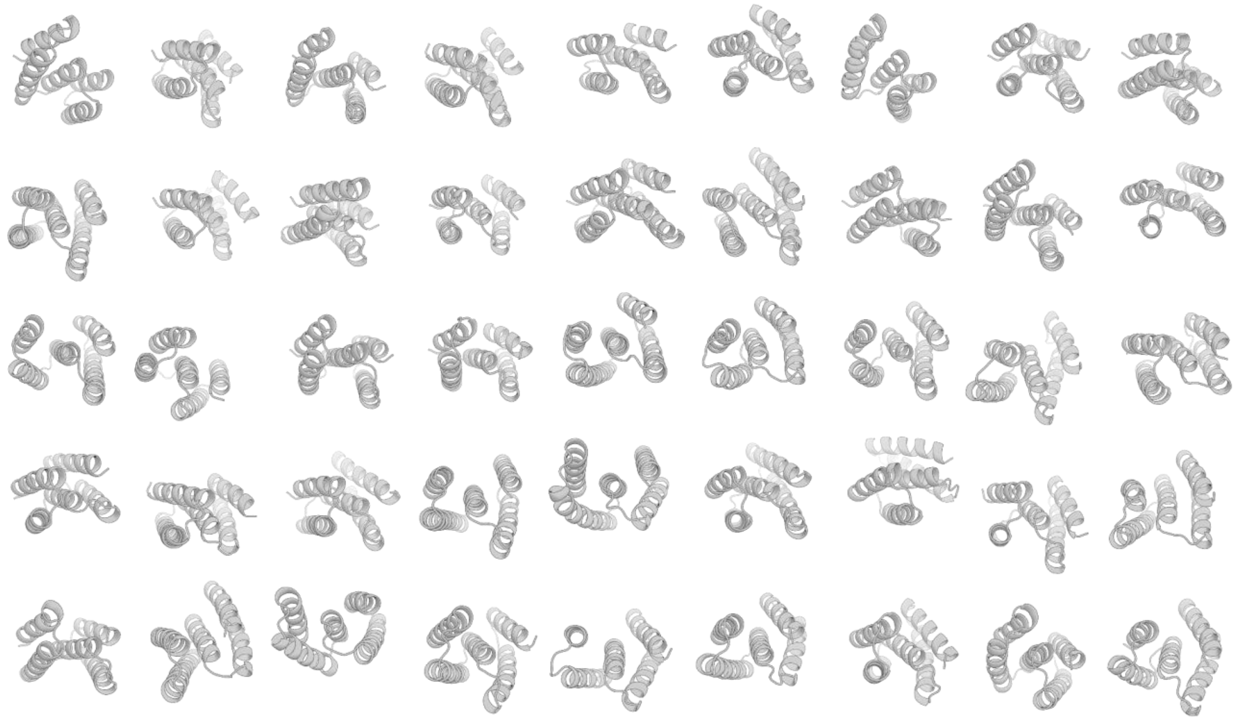

**Figure S1: Parametric groove scaffold library:** 45 scaffolds from the library of 18 thousand parametric groove scaffolds, demonstrating a range of supercoiling and helix distances to accommodate a range of helical peptide targets.

### Identification of weak binder hits from parametric designs in pilot experiment

The first helical peptide binder hits were identified in pilot experiments screening for binding using the nanoBiT split luciferase assay (methods). These kinetic binding experiments were performed in cell lysate with no control over protein concentration, so candidate binders were selected qualitatively for showing some increase in luminescence signal over time above background noise, indicating likely binding activity. Additional pilot experiments indicated that this binding activity was all at very weak affinities, likely >100 nM. Therefore, these initial candidates were not further characterized, but rather selected for additional design to yield higher affinity binders.

### Identification of  weak binders for NPY and GCG using extended parametric designs

We used the RF inpainting approach to extend the binding interfaces of NPY and GCG weak binders hits from parametric design. However, the characterized proteins displayed low affinity binding to their targets, which was not enough for diagnostic applications.

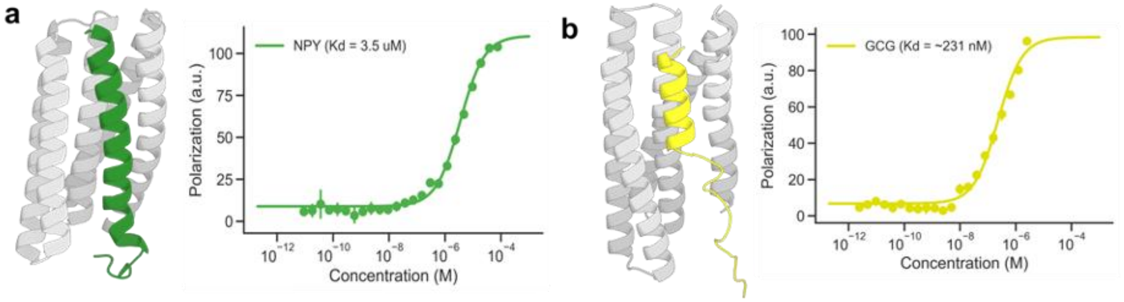


**Figure S2. Inpainted peptide binders bound  their targets with low affinity. (a)** NPY binder. **(b)** Glucagon binder. AF2 predictions of the proteins and peptides are shown on the left. FP binding data is shown on the right.

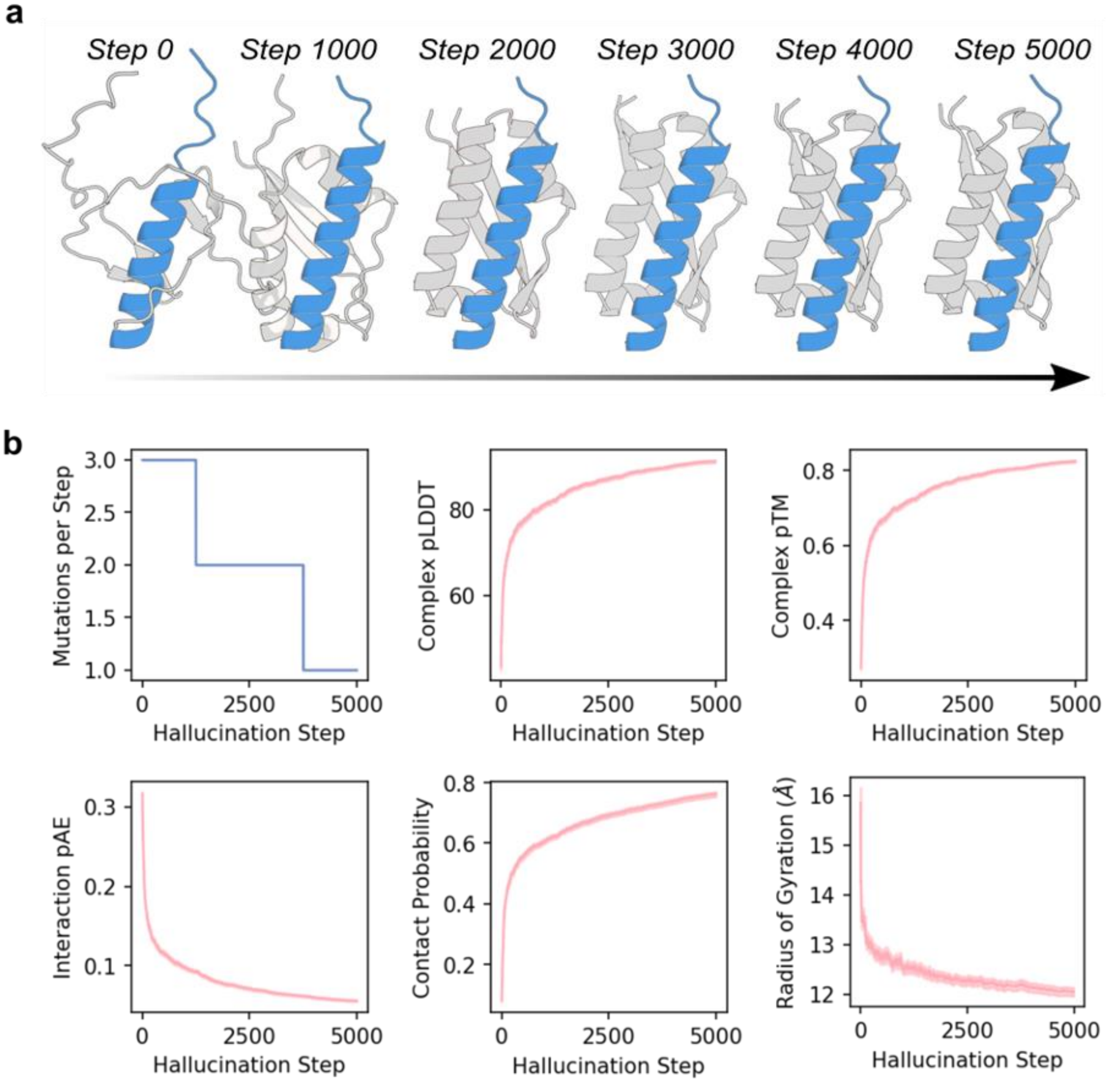


**Figure S3: Hallucinating Bid Binders with AlphaFold2: a)** Example hallucination trajectory generating 70 amino acid binders to the peptide Bid (blue). Initially, AlphaFold2 predicts an unstructured “binder”, but over 5000 steps, a binder is built up around the peptide. Crucially, no template structure is provided for the Bid peptide, allowing AF2 to predict its structure throughout. Note the predicted elongation of the helical structure in the peptide (blue, top) over the hallucination trajectory. **b)** Hallucination trajectories approximately converge after 5000 steps. Left to right, top to bottom: The mutation rate at each step is decayed throughout the trajectory (1250 x 3 steps, 2500 x 2 steps, 1250 x 1 step). More mutations initially helps speed up hallucination, while a lower rate later on allows more gradual refinement. The AF2 confidence (pLDDT, pTM) in the bound structure increases throughout trajectories, while the pAE between peptide and binder (known to be a good correlator of binding) decreases. The contact probability also trends to convergence over the trajectories, while the proteins typically become more compact (radius of gyration). N=96 trajectories.

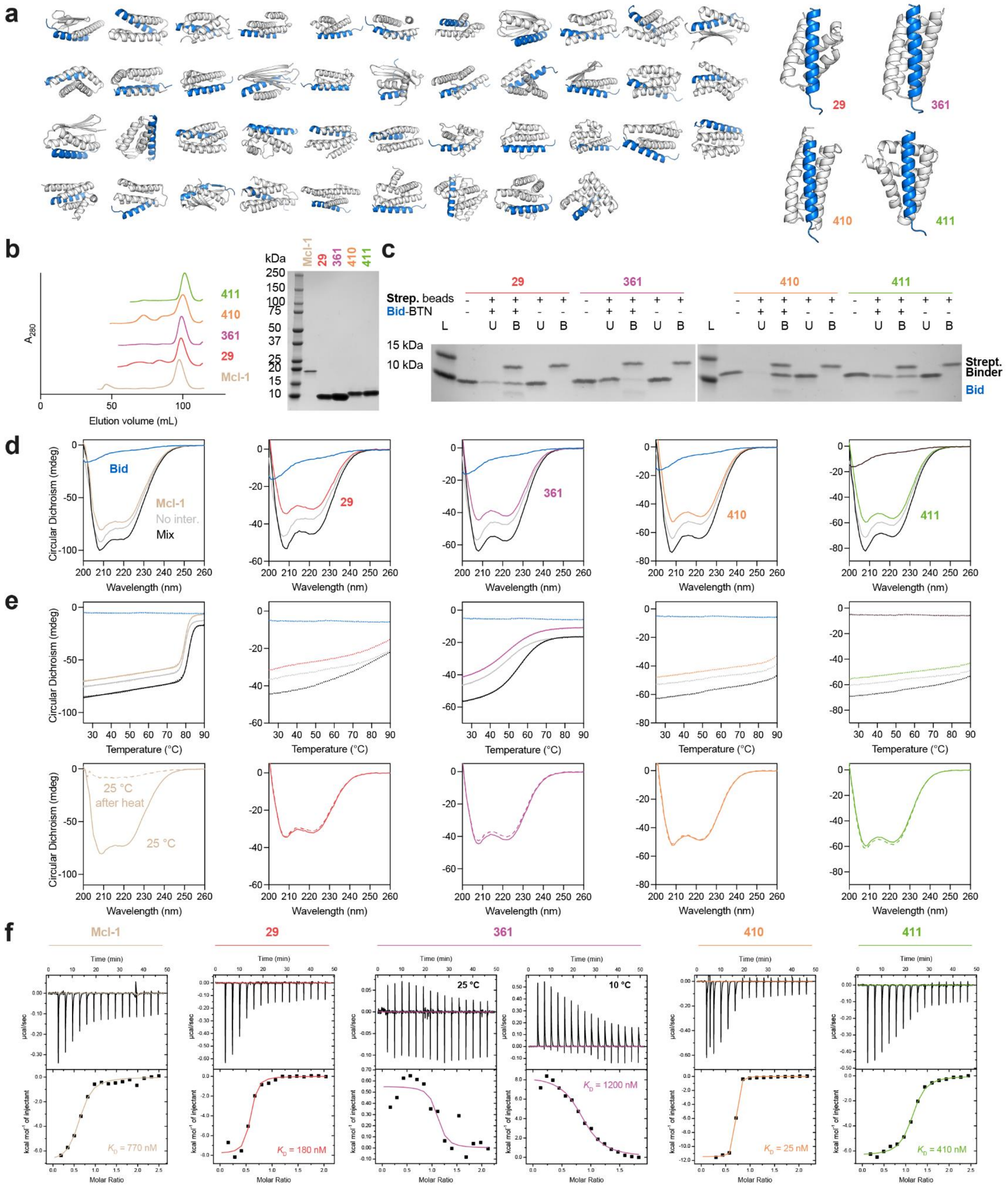


**Figure S4. Hallucinated Bid binders were stable and bound Bid peptide with high affinity. (a)** 47 hallucinated designs tested for initial experimental screening. **(b)** 4 designs were chosen for expression without Bid peptide. All expressed as monomeric proteins (assessed by preparative SEC) and were pure by SDS-PAGE. **(c)** All hallucinations could be pulled-down by biotinylated Bid immobilized on streptavidin magnetic beads. B = bound to bead, U = unbound, in supernatant. L = ladder. **(d)** Bid is unstructured in isolation by circular dichroism (CD), whereas all hallucinations were helical in isolation, as predicted from the hallucinated structure. A 1:1 molar ratio of binder:Bid (Mix) produced greater helical signal than that predicted by the isolated spectra (No inter.) suggesting binding is inducing helix formation. **(e)** Melting with CD showed that hallucinations were thermostable, and binding to Bid increased thermostability (where measurable). All hallucinations would remain folded, or refold after heating and cooling, in contract to the natural binder Mcl-1 which precipitated in the process. **(f)** ITC showed that hallucinations bound to Bid, with µM to nM $K_d$s.

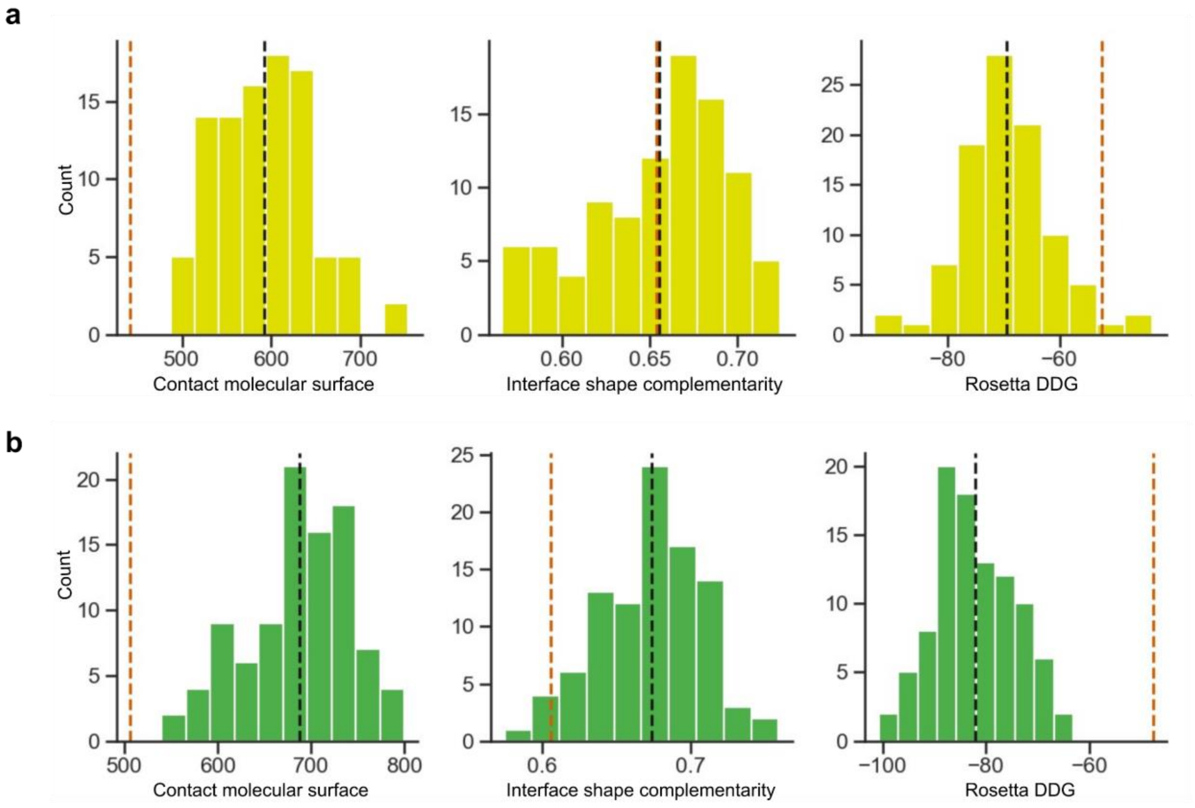


**Figure S5. Binding metrics for partially diffused binders. a)** Computational metrics for 96 ordered partially diffused glucagon binders showed significant improvement in contact molecular surface (a measure of interface size and quality) and Rosetta ddG (a measure of interface predicted energy) over the starting design (vertical red lines). Distribution means are shown in black. **b)** Computational metrics for 96 ordered partially diffused NPY binders showed significant improvement in contact molecular surface, Rosetta ddG, and interface shape complementarity (a measure of interface quality) over the starting design (vertical red lines). Means are shown in black.

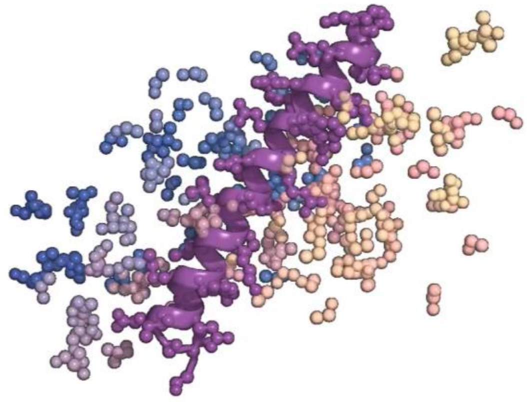

**Supplementary Video 1.** A video of the diffusion trajectory for the fully diffused PTH binder can be seen at https://www.bakerlab.org/wp-content/uploads/2022/11/diffusion_animation_PTHbinder_v6.mp4

| | GCG Binder | NPY Binder | PTH Binder | Bim Binder |
|---|---|---|---|---|
| **RMSD AF2 vs Design** | 0.62 Å | 0.61 Å | 0.78 Å | 0.80 Å |
| **AF2 interaction PAE** | 9.25 | 8.29 | 4.40 | 4.50 |
| **AF2 pLDDT for binder** | 95.52 | 93.41 | 94.3 | 96.6 |

**Table 1. Alphafold metrics for partially and fully diffused binders.**

**Parametric design of groove-shaped scaffold library and use for binder design**

The parametric groove-shaped scaffold library was sampled using a random sampling approach, where key parameters were selected randomly from distributions. An even distribution of bundle "lengths" was sampled, where each parametric helix was 15-19 residues long. A supercoiling value was randomly selected from a biased distribution favoring more supercoiled scaffolds, given these scaffolds were more likely to fail in the subsequent looping step. An average helix neighbor distance value was randomly selected from a normal distribution informed by native helical bundle geometries. The distance of each helix from its neighbors was independently randomly selected from a much tighter normal distribution centered at the preselected average helix neighbor distance value, to provide some noise within a given scaffold to helix distances and allow for heterogeneous amino acid selections. Values for helix phase and Z displacement were randomly sampled for each helix. The "groove" consisting of 3 helices was first sampled as a helical bundle using the Crick parameterization of alpha-helical coiled coils, around an imaginary central helix where the target was to later be docked. Next, the two buttressing helices were sampled with the same parameterization, but moved radially outward with randomly sampled helix neighbor distances as well as an additional randomly sampled tilt. This process was used to sample a set of 200k arrangements of 5 helices. Next, the Rosetta ConnectChainsMover was used to loop this set into approximately 135k successful scaffold backbones. These backbones were designed and filtered using Rosetta to yield a final library of 18 thousand scaffolds. This library was used to design binders to different helical peptide targets using an adapted version of the miniprotein binder design computational pipeline used by Cao *et al.*[5].

**Design of BIM peptide binders**

We also experimented with unconditional binder design for the apoptosis-related peptide Bim (DMRPEIWIAQELRRIGDEFNAYYARR; PDB: 6X8O)- providing RF*diffusion* only with the sequence and structures of the two peptides in helical conformations, and leaving the topology of the binding protein and the binding mode completely unspecified. From this minimal starting information, RF*diffusion* generated designs predicted by AF2 to fold and bind to the targets with high *in silico* success rates. We obtained synthetic genes encoding 96 designs for each target. Using yeast surface display, we found that 25 of the 96 designs bound to Bim (10nM, no avidity). The highest affinity design, which purified as a soluble monomer, bound too tightly for steady state estimates of the dissociation constant (Kd); global fitting of the association and dissociation kinetics suggest a Kd of ~100pM.  External potentials were used to promote interactions between the binder and target - specifically, the radius of gyration of the complex was minimized.
